# Graded regulation of microtubule-binding of Tau by the phosphorylation state of the proline-rich region in living neurons

**DOI:** 10.1101/2025.11.11.687775

**Authors:** Rinaho Nakata, Tomohiro Torii, Tomohiro Miyasaka, Hiroaki Misonou

## Abstract

Tau protein is a microtubule-associated protein that plays a crucial role in maintaining neuronal morphology and axonal transport. While phosphorylation is known to regulate Tau–microtubule interactions, the contribution of specific phosphorylation patterns *in situ* remains poorly understood due to the complexity of the intracellular environment. In this study, we combined fluorescence recovery after photobleaching (FRAP) in primary cultured rat hippocampal neurons with dephosphorylation-mimetic mutations and computational modeling to analyze the effects of phosphorylation on Tau-microtubule interaction. We particularly focused on the proline-rich region, of which phosphorylation has been studied in physiological and pathological perspectives, and generated a dephosphorylation-mimetic Tau mutant by substituting key phosphorylation sites with alanine residues and compared its microtubule-binding dynamics to those of WT-Tau in FRAP experiments. Experimental data, together with simulation-based parameter exploration, revealed that the overall number of phosphorylated sites plays a more dominant role than their specific locations in modulating Tau–microtubule affinity. These findings provide new insights into the post-translational regulation of Tau and establish a computational–experimental framework for interrogating intracellular protein dynamics.

## Introduction

Tau is a microtubule (MT)-associated protein (MAP) that, under normal physiological conditions, is predominantly localized to the axons of neurons (Binder et al., 1985; Weingarten et al., 1975). This has been further confirmed by recent studies using high-sensitivity detection methods, which show that normal Tau is exclusively localized to the axonal compartment *in vivo* (Kubo et al., 2019a). By promoting MT stabilization, Tau is thought to be involved in crucial neuronal processes, including axonal outgrowth, neuronal plasticity, and cargo transport (Caceres and Kosik, 1990; Dawson et al., 2010; Dixit et al., 2008; Drubin et al., 1985; Ebneth et al., 1998; Esmaeli-Azad et al., 1994; Hirokawa et al., 1988; Kempf et al., 1996; Liu et al., 1999; Samsonov et al., 2004; Trinczek et al., 1999). In neurodegenerative diseases known as tauopathies, including Alzheimer’s disease (AD), Tau mislocalizes to the soma and dendrites, where it ultimately forms neurofibrillary tangles (Grundke-Iqbal et al., 1986; Kosik et al., 1986). This cellular ‘mislocalization’ of Tau is considered a key step in the pathological cascade leading to neurodegeneration and synaptic dysfunction (Hoover et al., 2010; Ittner et al., 2010; McInnes et al., 2018). Indeed, studies in transgenic mouse models indicate that ectopic Tau expression leads to its abnormal somatodendritic distribution, a step that appears necessary for initiating the development of Tau pathology (Kubo et al., 2019b). Therefore, understanding how normal Tau is localized to and maintained within the axon, and why this mechanism fails in disease, is a critically important challenge (Wang and Mandelkow, 2015).

One key aspect of Tau localization is the MT-binding and its regulation by phosphorylation, as hyper-phosphorylated Tau is abundant in neurofibrillary tangles and loses MT-binding capability. While Tau is generally understood to bind MTs via its C-terminal MT-binding domain (MTBD), other regions such as the proline-rich regions (PRRs) are also known to contribute to this interaction (Goode and Feinstein, 1994; Gustke et al., 1994). More recently, structural analysis by cryo-electron microscopy has put forward a molecular model where the MT-binding region of Tau adopts an extended structure along the protofilament, stabilizing MTs by tethering tubulin dimers together (Kellogg et al., 2018). However, while these structural models provide a static snapshot of the binding, how this interaction is dynamically regulated within the complex environment of a living cell remains poorly understood.

A recent study has provided new perspectives on the roles of these domains in the proper axonal localization of Tau (Iwata et al., 2019). They demonstrated, surprisingly, that stable MT-binding via the MTBD is not essential for axonal localization; rather, excessively stable binding may lead to Tau’s mislocalization. Indeed, whereas a Tau mutant lacking the MTBD still localized properly to the axon, a dephosphorylation-mimetic mutant in the PRR2 region exhibited reinforced MT-binding and was mislocalized to the soma and dendrites. These findings strongly suggest that not only the MTBD, but also the PRR2 region and its phosphorylation state, are critically important for regulating the normal behavior of Tau within the neuron (Goode et al., 1997).

Although the importance of the PRR2 region has been suggested, the detailed molecular mechanism of ‘how’ and ‘to what quantitative extent’ its phosphorylation controls the binding dynamics of Tau and MTs remains largely unelucidated. One major issue is the difficulty of knowing and altering the phosphorylation state of endogenous Tau proteins in living neurons. Therefore, researchers have used phosphorylation- and dephosphorylation-mimetic mutations on exogenously expressed Tau using uncharged and negatively charged amino acid residues. However, it is still challenging to reliably monitor Tau mobility and MT-binding in situ, particularly in living neurons, and to relate the experimental data to kinetic parameters, such as association and dissociation rates. Conventional biochemical methods and FRAP analysis based on simple curve fitting have had limitations in capturing the full picture of this dynamic and quantitative regulation, calling for a more sophisticated approach (Mueller et al., 2010; Sprague and McNally, 2005).

To address these issues, we employed an integrated approach in this study, combining quantitative live-cell imaging (FRAP), site-directed mutagenesis, and computational modeling (Cowan and Loew, 2023). We systematically analyzed how the phosphorylation state of the PRR2 region affects the MT binding dynamics of Tau in living neurons. Here, we uncovered a novel mechanism wherein the phosphorylation state of the PRR2 region modulates its affinity for MTs in a graded manner. Furthermore, the development of our computational model enabled the estimation of the kinetic parameters underlying this regulation, providing a way for a deeper understanding of Tau’s functional control.

## Materials & Method

### Animals

This study utilized Sprague Dawley rats and Tau knockout mice (Tau-KO; (Dawson et al., 2001; Harada et al., 1994)). All animal procedures were approved by the Institutional Animal Care and Use Committee of Doshisha University.

### Neuronal cultures

Primary hippocampal neurons were prepared from embryonic day 17–18 Sprague Dawley rats, following the previously described method (Misonou et al., 2008). Dissected hippocampi were treated with 0.25% trypsin for 15 minutes at 37°C, followed by dissociation in poly-L-lysine-coated coverslips at a density of 40,000 cells per slip. Neurons were cultured in 5% CO₂ at 37°C and on day 4 in vitro (DIV), Ara-C (final concentration: 5 µM) was added to inhibit glial cell proliferation. Some aliquots were frozen using the DMSO-based method (Ishizuka and Bramham, 2020). Briefly, neurons were suspended in a freezing solution containing 20% DMSO and 80% FBS, then transferred into cryotubes and stored in liquid nitrogen. Prior to plating, frozen neurons were thawed, suspended in pre-warmed culture medium, counted, and plated at a density of 40,000 cells per coverslip. After an initial 4–6 h incubation to allow for attachment, culture medium was replaced with conditioned medium from astrocyte cultures (Misonou and Trimmer, 2005).

### DNA Constructs

Human Tau cDNA corresponding to the 0N4R isoform (1–383 amino acids) was obtained from pLVSIN-Tet3G-AcGFP-His-Tau0N4R, which was generated as described previously (Iwata et al., 2019).Mutations were introduced into the proline-rich region (PRR2; amino acids 140–185) and the microtubule-binding domain (MTBD; amino acids 186–309) by PCR-based site-directed mutagenesis using PrimeSTAR Max DNA Polymerase (Takara) or a KOD -Plus- Mutagenesis Kit (TOYOBO), with the WT construct as a template.

All constructs express human 0N4R Tau fused to AcGFP at the N-terminus and a His tag at the C-terminus under the control of a TRE3G promoter in the pLVSIN-Tet3G lentiviral backbone.

### Lentiviral Vector Production

Lentiviral vectors were produced using the calcium phosphate method (Chen and Okayama, 1987). Human embryonic kidney (HEK) 293T cells were cultured in DMEM supplemented with 10% fetal bovine serum. Cells at 70% confluency were transfected with 12 μg of pLVSIN containing GFP-Tau, 6 μg of packaging plasmids (pRSV-Rev, pMD2.G, and pMDLg/pRRE). After incubation, the medium was replaced, and lentiviral particles were harvested 48 hours post-transfection. The viral solution was filtered and stored at –80°C until use. Lentiviral titers were determined using the Lenti-X qRT-PCR Titration Kit (Takara).

### Lentiviral transduction and induction of protein expression

Primary hippocampal neurons were transduced with pLVSIN-Tet3G-AcGFP-His-Tau0N4R (WT or mutant) lentiviral particles at the day of plating at a multiplicity of infection (MOI) of 10–50. After overnight incubation, neurons on coverslips were transferred to new six-well plates containing 1–1.5 mL of astrocyte-conditioned medium prepared as described previously (Misonou and Trimmer, 2005).

For Tau induction, the medium was replaced with fresh conditioned medium containing 1 µg/mL doxycycline (Dox) for 1 h, followed by transfer back to the original six-well plates with astrocytes, as described previously (Iwata et al., 2019). Expression of AcGFP-tagged Tau (both WT and mutant) was confirmed by fluorescence. For AcGFP–α tubulin induction, neurons were treated with 1 µg/mL Dox for 8 h prior to live-cell imaging, and Dox was maintained in the culture medium during imaging to sustain expression.

### Immunofluorescence staining

Neurons were fixed in 4% paraformaldehyde in PBS for 20 min at room temperature. Blocking and permeabilization were performed in 4% dried milk and 0.1% Triton X-100 in TRIS-buffered saline (TBS) for 30 min. Primary and secondary antibodies were diluted in the same buffer and applied in separate incubation steps of 1 h and 45 min, respectively. Coverslips were mounted on glass microscope slides using ProLong Diamond antifade reagent (Thermo Fisher Scientific). For immunostaining, neurons were incubated with anti-MAP2 antibody (clone AP20, 1:1000, Sigma-Aldrich) followed Alexa Fluor 594-conjugated anti-mouse IgG secondary antibody (1:2000, Thermo Fisher Scientific).

### Live-cell imaging (FRAP)

Live imaging experiments were performed using a Leica TCS SP8 confocal microscope(Leica Microsystems, Wetzlar, Germany) equipped with a HC PL APO 63×/1.40 OIL CS2 objective lens and HyD detectors. Neurons expressing GFP-fused Tau (either WT or mutant) were maintained at room temperature (approximately 25–27°C) during imaging. After identifying the region of interest (ROI), the fluorescence within a circular ROI (3–5 µm in diameter) was photobleached by brief exposure (0.5–1 s) to the 488 nm laser at an intensity sufficient to induce bleaching. Recovery of fluorescence was recorded at 1-second intervals for up to 70 s post-bleaching. Images were processed using ImageJ (National Institutes of Health), subtracting background signals and correcting for ongoing bleaching due to excitation. Fluorescence values were then normalized so that the minimum and maximum intensity were set to 0 and 1, respectively (Jensen et al., 2017).

### Statistical analysis

All statistical analyses were performed on GraphPad Prism software (GraphPad Software, San Diego, CA). The recovery phase from the minimum was fitted with the one-phase association model: Y=Y_0_ + (plateau-Y_o_) × (1-10^-k×X^). The difference of the fitted curves among different Tau variants or in different areas in neurons was analyzed with an F test for the null hypothesis that a curve with shared parameters (slope k and plateau) fits better than those with different parameters for each data set. We also fitted data from individual neurons and obtained an averaged slope and plateau for each mutant.

### Virtual Cell Simulation

To further elucidate the kinetics of Tau diffusion and binding, we constructed a reaction-diffusion model using Virtual Cell (https://vcell.org/). This model integrates experimental FRAP conditions—namely, 1) diffusion of free Tau species,2) reversible Tau–MT binding, and (3) localized photobleaching—within a three-dimensional neuronal compartment. Key aspects of the modeling approach are detailed below.

#### Model Geometry and Initial Conditions

- 3D Axon Geometry: We defined a cylindrical-like geometry (length = 200 µm, diameter = 2 µm) to approximate the axon.
- Initial conditions: we used the homogeneous single compartment simulation with estimated concentrations (shown in Table 2) and kinetics to establish the initial condition with the axonal geometry.
- MT Concentration: The microtubule concentration (MT) was set between 45 and 50 µM in the cytosol, reflecting an effective tubulin/Tau-binding capacity estimated from in vitro data. These biochemical data were obtained from previously published studies(Iwata et al., 2019).

##### Reaction Kinetics

All reactions were modeled using mass-action kinetics. The primary reactions included:

1. Tau–MT Binding

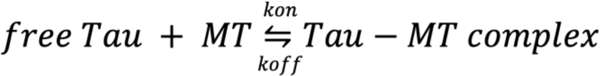

- Association rate constant? Forward rate constant (Kon): Typically set to 1.0 s⁻ ¹·µM⁻ ¹, based on previous Tau-binding studies.
- Dissociation rate constant? Reverse rate constant (Koff): Varied between 0.22 and 4.0 s⁻ ¹ in different simulation runs to explore how altered off-rate affect FRAP recovery profiles.
2. Photobleaching Reactions A bleaching reaction was applied to convert fluorescent Tau (exfTau, enfTau, etc.) to nonfluorescent forms (exTau, enTau, etc.) within the chosen bleaching region. In the Virtual Cell model, this was implemented as:

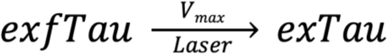

- bleaching Region and Duration: The laser (represented as the species Laser) was defined to exist only in a rectangular zone from x>100 μmx > 100 uM to x<104 μmx for 0.5 s at t≈10s of the simulation.
- Bleach Rate (Vmax): Typically set to 50 s⁻ ¹·µM⁻ ¹, ensuring rapid conversion of fluorescent Tau to its bleached counterpart within the specified ROI.
- All species and used in the simulation are listed in Table 1.
3. Diffusion Each species (free Tau, bound Tau, bleached Tau, etc.) was assigned a diffusion coefficient:

- Free Tau (e.g., exfTau, enfTau): 10.0 µm²/s (indicative of cytoplasmic diffusion for a 50–60 kDa protein).
- Bound Tau (e.g., exfTaub, enfTaub): 0.001 µm²/s, reflecting that Tau complexed with MTs is effectively immobile.
- Bleached Tau (e.g., exTau, enTau): Same diffusion coefficient as free Tau, assuming bleaching does not alter diffusion but only fluorescence.

**Table 1.** Definitions and Roles of Species in the Simulation Model.

| Species Name | Description | Role |
| --- | --- | --- |
| MT |  | Microtubule |
| Laser |  | Light source for bleaching |
| enTau |  | Endogenous Tau (bleached) |
| enfTau |  | Endogenous fluorescent Tau |
| enTau<sub>b</sub> |  | MT-bound endogenous Tau (bleached) |
| enfTau<sub>b</sub> |  | MT-bound endogenous fluorescent Tau |
| fast_exTau |  | Fast-binding exogenous Tau (bleached) |
| fast_exfTau |  | Fast-binding exogenous fluorescent Tau |
| fast_exTau<sub>b</sub> |  | MT-bound fast- binding Tau (bleached) |
| fast_exfTau<sub>b</sub> |  | MT-bound fast-binding fluorescent Tau |
| slow_exTau |  | Slow-binding exogenous Tau (bleached) |
| slow_exfTau |  | Slow- binding exogenous fluorescent Tau |
| slow_exTau<sub>b</sub> |  | MT-bound slow-binding Tau (bleached) |
| slow_exfTau<sub>b</sub> |  | MT-bound slow-binding fluorescent Tau |

**Table 2.** Parameter Estimation Based on Biological Date.

|  | *Initial concentrations* | *Basis for Estimation* |
| --- | --- | --- |
| MT | 50 µM | Quantitative Western Blotting |
| Endogenous Tau | 5 µM | Quantitatieed from immunofluorescence images <sup>a)</sup> |
| Exogenous Tau | 30 µM | Quantitatieed from immunofluorescence images <sup>a)</sup> |
| *Diffusion* |  |  |
| MT | 0.001 µm<sup>2</sup>/s | Virtually immobile |
| Unbound Tau species | 10 µm<sup>2</sup>/s | FRAP recovery curves |

a) Values derived from lwata et al., (2019).

##### Numerical Setup

- Partial Differential Equations (PDEs): Each species was governed by a PDE comprising a reaction term (binding/unbinding, bleaching) and a diffusion term.
- Mesh and Time Steps: For the 3D simulations, a spatial mesh of 401×5×5×5 elements was typically used. Time steps were solved with the Sundials Stiff PDE Solver under variable-step settings (default time step = 0.05 s, max time step = 0.1 s).
- Boundary Conditions: Zero-flux boundaries were assigned at the cylinder edges (i.e., no Tau exchange outside the domain), while the extracellular compartment was included for completeness.

##### Simulation Protocol

Simulations were run from t=0 to t=80 s. The bleaching step was triggered between t = 10 and t=10.5 s, at which point fluorescent Tau species in the designated region were rapidly converted to the bleached state. Post-bleach fluorescence recovery was monitored by measuring the re-entry of fluorescent Tau from adjacent regions and binding/unbinding events. Multiple runs were performed:

- Baseline: k_off_ = 3.0s^-1^, free Tau = 5 µM, etc.
- Parameter Sweeps: k_off_ was varied (e.g., 0.22, 0.7, 2.5, 4.0) to capture different binding affinities, and the corresponding FRAP curves were compared to experimental data.

##### Data Analysis and Comparison with FRAP

The simulated fluorescence intensity profiles were mapped to a normalized scale analogous to experimental ImageJ processing. Recovery curves from the Virtual Cell model were then overlapped with measured FRAP data to evaluate how well different parameter sets (diffusion coefficients, on/off rates) recapitulated the observed Tau kinetics. Overall, these simulations reinforced the biphasic recovery noted experimentally and helped pinpoint the kinetic parameters most crucial for describing Tau’s rapid phase (diffusion-dominated) and slower phase (MT-binding-dominated) recovery.

## Result

To investigate the dynamics of Tau protein interaction with MTs in living neurons, we employed Fluorescence Recovery After Photobleaching (FRAP) microscopy. We assumed that Tau protein dynamically shuttles between a freely diffusing state (Diffusion-Tau) and a state bound to MTs (Binding-Tau) (Figure 1A). Therefore, FRAP data would provide the relative contributions of these states for further analysis (Figure 1C).

**Figure 1.**
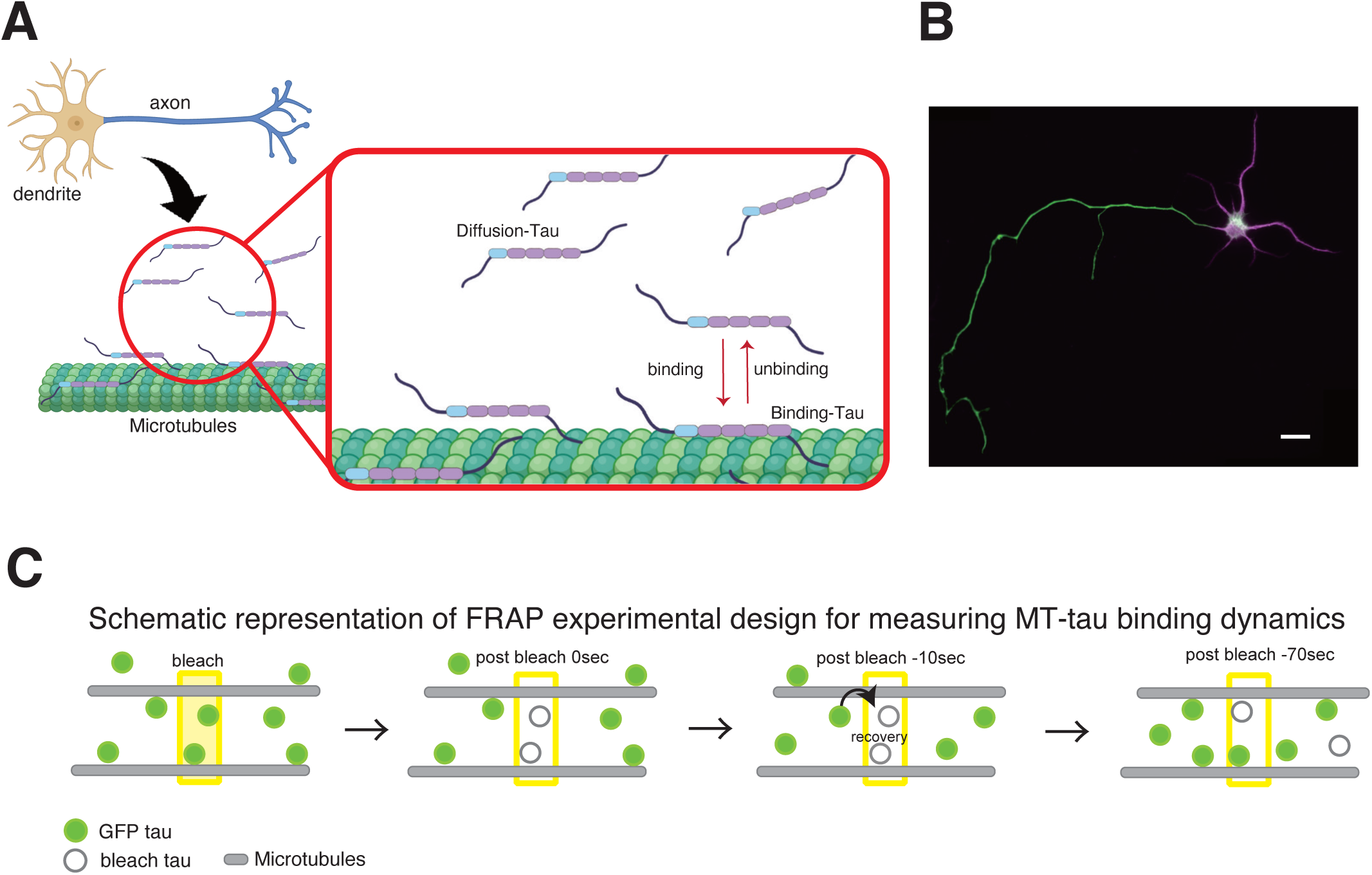
FRAP analyses of GFP-tagged Tau in wild-type and Tau-knockout (TKO) mice neurons. **(A)** Schematic diagram illustrating how Tau protein dynamically shuttles between a freely diffusing state (Diffusion-Tau) and a state bound to microtubules (Binding-Tau) within the axon. Schematic was created with Biorender.com. **(B)** A representative image of a primary cultured rat hippocampal neuron (DIV 4-7) expressing GFP-tagged WT-Tau. Tau is shown to be enriched in the axon, where FRAP experiments were performed. Scale bar, 20 µm. **(C)** Schematic illustration of FRAP. FRAP would be dominated by diffusion and MT-binding of Tau molecules.

### The Phosphorylation State of the PRR2 Domain Bidirectionally Regulates Tau’s MT-Binding Dynamics

To begin our investigation, we first characterized the dynamics of wild-type GFP-tagged Tau (WT-Tau) expressed in primary cultured hippocampal neurons (Figure 1B). For these studies, we used immature neurons at DIV 4-7. This period corresponds to developmental stages 3 to early 4, a critical window for axonal specification and the establishment of neuronal polarity. In immature neurons at this stage, the dynamic reorganization of MTs is active; for instance, acetylated tubulin, a marker for stable MTs, has been shown to accumulate in the distal axon (Hammond et al., 2010). In parallel with these changes in MT dynamics, the process of ‘axonal localization’, wherein Tau is selectively transported to and enriched in the axon, is actively underway (Figure 1B). Therefore, we considered DIV 4-7 to be the optimal stage to elucidate the fundamental binding mechanisms between Tau and MTs that underlie its axonal localization. The expression of GFP-Tau in our inducible-expression system was approximately 2-fold higher than that of endogenous Tau, as previously quantified and reported in the same experimental system (Iwata et al., 2019).

We have previously analyzed expressed Tau using FARP and showed that GFP-Tau exhibited dynamics largely dependent on MT-binding (Iwata et al., 2019). We were able to replicate these results in this study (Figure 2B). Therefore, we next investigated how the phosphorylation state of the proline-rich region 2 (PRR2) modulates Tau’s interaction with MTs. Phosphorylation within this domain is known to reduce Tau’s affinity for MTs and is implicated in Alzheimer’s disease pathology (Kiris et al., 2011; Schwalbe et al., 2015). Indeed, it has been previously reported that a phospho-mimetic mutant, where eight key sites in PRR2 (Ser198, Ser199, Ser202, Ser205, Ser212, Thr214, Thr231, and Ser235; Figure 3A) were substituted with glutamate (8E-Tau), exhibits significantly faster FRAP recovery than WT-Tau, consistent with a more transient, weakly-bound state (Iwata et al., 2019; Rodríguez-Martín et al., 2013). Furthermore, we and others have shown that dephosphorylation-mimetic mutants with alanine substitution (8A-Tau) result in reduced FRAP, indicating more stable binding of Tau to MTs ( Rodríguez-Martín et al., 2013; Iwata et al., 2019).

**Figure 2.**
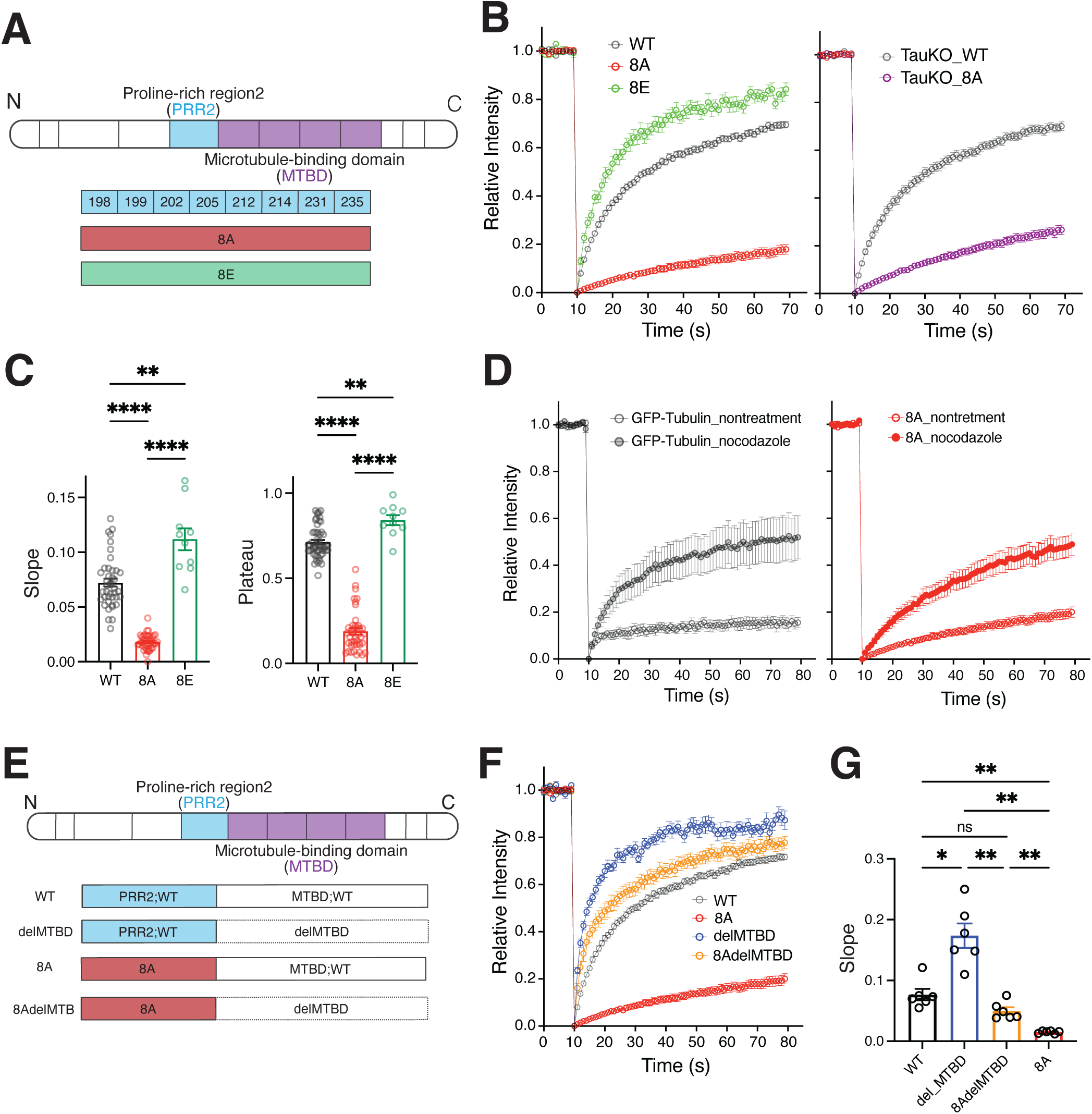
FRAP analyses of dephosphorylation-mimetic mutant of Tau (8A-Tau) in the proline-rich region 2 (PRR2) in Tau in wild-type and TKO mice neurons. **(A)** Schematic of WT-Tau and mutants mimicking dephosphorylated (8A) and phosphorylated (8E) states. The 8A and 8E mutants contain alanine or glutamate substitutions, respectively, at eight key phosphorylation sites: S198, S199, S202, S205, S212, T214, T231, and S235. **(B)** FRAP curves. Left, wild-type rat primary neurons comparing WT-Tau, dephospho-mimetic 8A-Tau, and phospho-mimetic 8E-Tau; 8A-Tau shows markedly slower recovery, whereas 8E-Tau recovers faster than WT-Tau. The right panel compares 8A-Tau dynamics in wild-type versus Tau-knockout (TKO) mice neurons, showing the effect persists in the absence of endogenous Tau. **(C)** Quantification of the recovery slope (left) and plateau (right) for WT, 8E-Tau, and, 8A-Tau in rat neurons. **(D)** FRAP curves for GFP□α□tubulin (left) and 8A-Tau (right) with or without treatment with the microtubule-depolymerizing agent nocodazole in rat neurons. Nocodazole treatment confirmed that the measured dynamics are MT-dependent. **(E)** Schematic of Tau deletion constructs: a mutant lacking the microtubule-binding domain (delMTBD) and a double mutant combining the 8A mutation with the MTBD deletion (8AdelMTBD). **(F)** Representative FRAP curves for the constructs shown in (E). Deletion of the MTBD results in rapid and complete fluorescence recovery, regardless of the 8A mutation. **(G)** Quantification of the FRAP plateau values for the constructs in (F). The low mobile fraction phenotype of 8A-Tau is completely rescued by the deletion of the MTBD. Data are presented as mean ± SEM (WT, 0.0766 ± 0.0094; delMTBD, 0.1739 ± 0.0201; 8AdelMTBD, 0.0499 ± 0.0059; 8A, 0.0150 ± 0.0009; n = 6 for all groups). Statistical analysis was performed using Games-Howell’s multiple comparisons test (**p < 0.01; ns, not significant).

**Figure 3.**
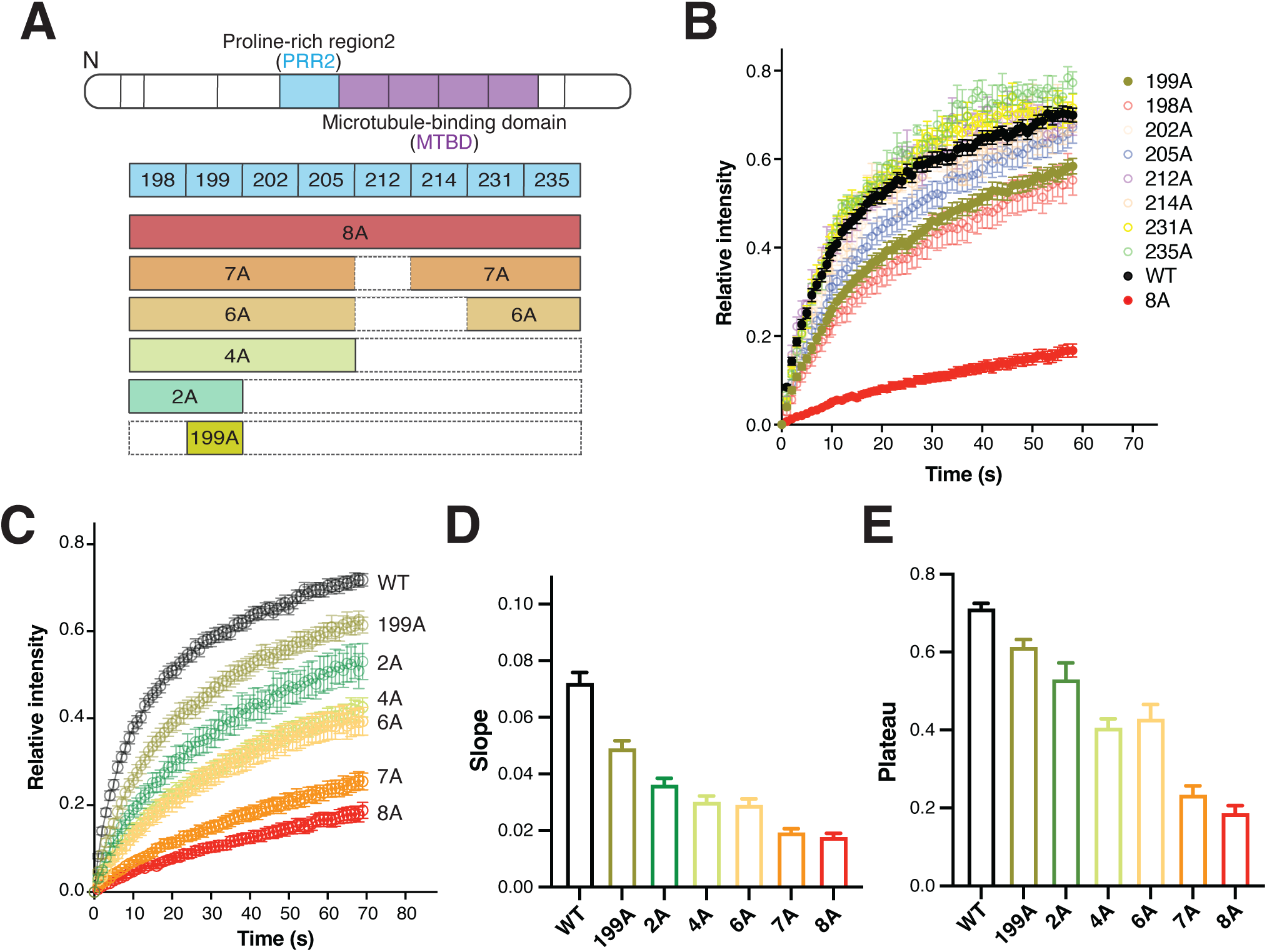
FRAP analyses of dephosphorylation-mimetic mutants with varying number of alanine substitution in rat primary neurons. **(A)** Diagram showing the six mutants used in the study. **(B)** FRAP recovery curves (mean ± SEM) for single□Ala mutants at PRR2 sites compared with WT in primary hippocampal neurons from rat. Quantification of the recovery slope and plateau for WT and single-alanine Tau mutants (198A and 199A) expressed in rat primary neurons. Compared with WT, a significant reduction in both recovery slope and plateau was observed not only in the 8A mutant but also in the single-alanine mutants 198A and 199A, whereas all other single-site substitutions showed no significant difference.Data are presented as mean ± SEM (slope: 0.0723 ± 0.0038 for WT, n = 41; 0.0565 ± 0.0052 for 198A, n = 20; 0.0522 ± 0.0047 for 199A, n = 40; plateau: 0.712 ± 0.019 for WT, n = 56; 0.586 ± 0.026 for 198A, n = 20; 0.584 ± 0.022 for 199A, n = 33. Welch’s ANOVA followed by Games–Howell’s test; plateau—198A p < 0.0001, 199A p < 0.0001; slope—198A p = 0.0023, 199A p = 0.0053; **** p < 0.0001; ** p < 0.01). **(C)** FRAP data of all six mutants with WT-Tau. Mean ± SEM from multiple cells are shown. **(D)** Slope obtained from individual cells. **(E)** Plateau obtained from individual cells.

We have been able to reproduce these results in that 8E- and 8A-Tau exhibited remarkably opposite effects on Tau FRAP: 8E-Tau showed significantly increased slope and plateau, while FRAP of 8A-Tau was greatly reduced, as compared to WT-Tau (Figure 2B), in curve fitting with single exponential functions. Statistical comparisons of slope and plateau revealed significant differences (Figure 2C): For slope, Welch’s ANOVA (2.000, 16.47) = 120.3, p < 0.0001 followed by Games-Howell’s multiple comparisons test (WT vs. 8A, p < 0.0001; WT vs. 8E, p = 0.0458; 8A vs. 8E, p = 0.0004), and for plateau, Welch’s ANOVA (2.000, 25.61) = 316.8, p < 0.0001 followed by Games-Howell’s multiple comparisons test (WT vs. 8A, p < 0.0001; WT vs. 8E, p = 0.0038; 8A vs. 8E, p < 0.0001). We also performed the FRAP experiments in Tau-KO neurons to test whether the presence of endogenous Tau affects the Tau behavior. We obtained similar WT FRAP data in these Tau-KO neurons with no discernable differences in the recovery kinetics (Figure 2B). Also, quantitative analysis confirmed that the recovery rate constant (K) was significantly reduced for 8A-Tau compared to WT-Tau (WT: 0.085±0.011 vs. 8A-Tau: 0.016±0.001; mean ± SEM, n= 6 for both; unpaired t-test, t (10) = 6.342, p<0.0001). Because the difference was more substantial between WT- and 8A-Tau than with 8E-Tau and would provide a large window to analyze the effects of individual phosphorylation sites, we chose 8A-Tau as the template for the rest of the study here.

Next, we treated 8A-Tau expressing rat neurons with nocodazole. This treatment led to a significant acceleration of fluorescence recovery of GFP-α-tubulin, which was not robust before the treatment, suggesting the partial disruption of MTs by the drug (Figure 2D). Nocodazole also resulted in a significant augmentation of recovery of 8A-Tau (Figure 2D). Together with the result of Tau-KO neurons, we concluded that the slow dynamics of 8A-Tau is not due to aggregation from overexpression but rather a strong interaction with MTs. Also, the fact that the 8A and 8E mutations shift the dynamics in opposite directions relative to WT (Iwata et al., 2019; Rodríguez-Martín et al., 2013) provides strong evidence that this reflects a specific regulatory mechanism of MT-binding by the phosphorylation state, rather than a non-specific artifact of amino acid substitution.

### The PRR2 Domain Modulates Tau-MT Interactions Independently of the Canonical MTBD

We next sought to determine whether the pronounced effect of the 8A dephospho-mimetic mutation in PRR2 is dependent on the canonical MT-binding domain (MTBD). For this purpose, we generated and analyzed several GFP-tagged Tau constructs: WT-Tau, 8A-Tau, delMTBD-Tau (PRR2-WT, MTBD deleted), and a double mutant, 8AdelMTBD-Tau (PRR2-8A, MTBD deleted) (Figure 2E). As shown previously, 8A-Tau exhibited very slow recovery. In contrast, delMTBD-Tau showed substantially faster recovery (Figure 2F), indicating, as expected and previously reported (Rodríguez-Martín et al., 2013), that the MTBD is a primary contributor to Tau’s interaction with MTs. When the 8A mutation in PRR2 was combined with the deletion of the MTBD (8AdelMTBD-Tau), the FRAP dynamics were significantly altered compared to delMTBD-Tau alone. Specifically, 8AdelMTBD-Tau exhibited a markedly slower recovery rate than delMTBD-Tau (Figure 2F). Quantitative analysis of the recovery rate constant (K) corroborated these findings, showing the K value for delMTBD-Tau was significantly higher than that for 8AdelMTBD-Tau (Figure 2G) (delMTBD-Tau: 0.174±0.020 vs. 8AdelMTBD-Tau: 0.050±0.006, mean ± SEM; p=0.0028; Games-Howell’s multiple comparisons test). Similarly, the recovery plateau was evaluated. While 8AdelMTBD-Tau showed a tendency for a lower plateau compared to delMTBD-Tau, this difference was not statistically significant (delMTBD-Tau: 0.873±0.039 vs. 8AdelMTBD-Tau: 0.778±0.026, mean ± SEM; p= 0.2206; Games-Howell’s multiple comparisons test). These results collectively suggest that the 8A dephospho-mimetic mutation in the PRR2 domain participates in MT-binding independently but also enhances Tau’s interaction with MTs through MTBD.

### The Number of Dephospho-mimetic Sites in PRR2 Gradually Modulates Tau-MT Binding

Based on the observation that 8A-Tau exhibited significantly altered MT-binding dynamics, we sought to determine the responsible site(s) among the eight PRR2 phosphorylation sites by generating and testing a series of single alanine mutants. However, we did not detect substantial changes with any of the single mutants (Figure 3B), with a few exceptions including 199A single substitution mutant, which exhibited significantly lower recovery than WT-Tau (Figure 3A and 3B). This prompted us to consider whether the number of mutated sites within the PRR2 region, rather than a few dominant sites, contributes to this effect in a graded manner. To test this, we generated additional GFP-tagged Tau mutants with seven (7A-Tau) or six (6A-Tau) alanine substitutions within the same set of sites (Figure 3A). We also generated a few 2A and 4A mutants with selected sites out of all the combinations of 2As and 4As (Figure 3A). We picked 198A/199A (2A) and 198A/199A/202A/205A (4A) because the 199A single mutant exhibited the most significant effect on FRAP within the eight single mutants (Figure3C).

Interestingly, we observed somewhat gradual and progressive changes in Tau FRAP with increasing number of alanine substitution (Figure 3C), although the effects were seemingly nonlinear. WT-Tau showed the fastest recovery, while from 199A single mutant to 8A-Tau the recovery gradually became slower (Figure 3C). Quantitative analysis of the FRAP recovery rate constant (K) revealed significant differences among the groups (Figure 3D) (Welch’s ANOVA, W (3.000, 25.43) = 20.41, p< 0.0001). As expected, WT-Tau exhibited the highest recovery rate constant (Figure3D) (mean K= 0.081±0.007, n= 12; mean ± SEM), which was significantly higher than all dephospho-mimetic mutants analyzed (p≤0.0018 for all comparisons against WT; Games-Howell’s post hoc test). As shown in Figure 3D, a clear trend of decreasing mean K values was observed from 199A single mutant to 8A-Tau, although not all pairwise comparisons reached the significance level (see Figure legend). Similarly, the plateau showed graded reduction with the increasing number of alanine substitutions (Figure 3E).

Taken together, these results demonstrate that the number of dephospho-mimetic sites within the PRR2 region gradually modulates Tau’s binding dynamics with MTs. This supports the hypothesis that the overall number of non-phosphorylated residues in PRR2, rather than the status of a few critical sites impacts Tau’s affinity for MTs in living neurons.

### Computational Modeling of Tau-MT Dynamics Using Virtual Cell

To gain deeper insights into the mechanisms of Tau-MT interaction, we developed a computational model using the Virtual Cell (VCell) platform. A key motivation for our approach was to use biologically realistic parameters as model constraints and estimate reaction kinetics of Tau-MT interaction from the model. We determined key parameters (constraints) such as initial protein concentrations and diffusion coefficients directly either from our experimental data, literature, or empirical deduction (Table 2). The initial concentrations of proteins for the simulation were estimated as follows. The concentration of microtubules (MTs) was determined based on structural estimations from literature values and quantitative immunoblotting(Hagita et al., 2021; Yonemura et al., 2023). The concentration of endogenous Tau (enTau) was estimated with reference to studies establishing immunofluorescence as a reliable method for assessing endogenous protein distribution(Kubo et al., 2019a). The expression level of exogenous Tau (exTau) was subsequently determined as a ratio relative to enTau by comparing the fluorescence intensities of transfected versus non-transfected neurons, an approach consistent with previous work that quantified such expression ratios (Iwata et al., 2019).

To establish the diffusion coefficient for Tau, we used FRAP data from delMTBD-Tau, which we would diffuse freely within the cytoplasm without significant microtubule interaction. Virtual FRAP experiments were performed in VCell within a simple cylindrical geometry representing an axon. We then iteratively ran the simulation, varying the diffusion coefficient (D), and the best-fit value for D was determined by finding the value that minimized the sum of squared residuals between the simulated and experimental curves. This procedure yielded an estimated diffusion coefficient of approximately 12.5 µm²/s for ΔMTBD-Tau, which is consistent with a protein of its size diffusing in the cytoplasm. Based on this empirically derived value, we set the diffusion coefficient for free, WT-Tau in our model to 10.0 µm²/s. The diffusion coefficient of Bound Tau (e.g., exfTaub, enfTaub) was set to 0.001 µm²/s, considering that on the FRAP timescale (∼1 min), Tau bound to microtubules, which showed negligible FRAP (Figure 2D), could be treated as virtually immobile.

Using these constraining parameters, we first tested a simple "single-state model" (Figure 4A), where Tau interacts with MTs with a single set of on and off binding kinetics (Table 3). By monitoring the residuals and goodness of fit (Figure 4D), we approximated K_on_ and K_off_ values and the associated K_D_ to fit simulation data with experimental data (Figure 4B and 4C). We obtained the best fit value with a K_D_ of 3 µM (Figure 4C). Considering that our Tau protein includes physiological phosphorylation states in developing neurons, this value was in good agreement with the wide range of K_D_ values (∼0.01 µM to ∼2 µM) reported in the literature for recombinant Tau(Butner and Kirschner, 1991; Fauquant et al., 2011; Hervy and Bicout, 2019; Makrides et al., 2004). Since the simulation is based on the advection-diffusion-reaction equation, reactions faster than certain rate constants should be invariable. We are only able to say that the simulation failed with rate constants smaller than 0.3 µM□¹s□¹ (for k_on_) and 0.1 s□¹ (for k_off_).

**Figure 4.**
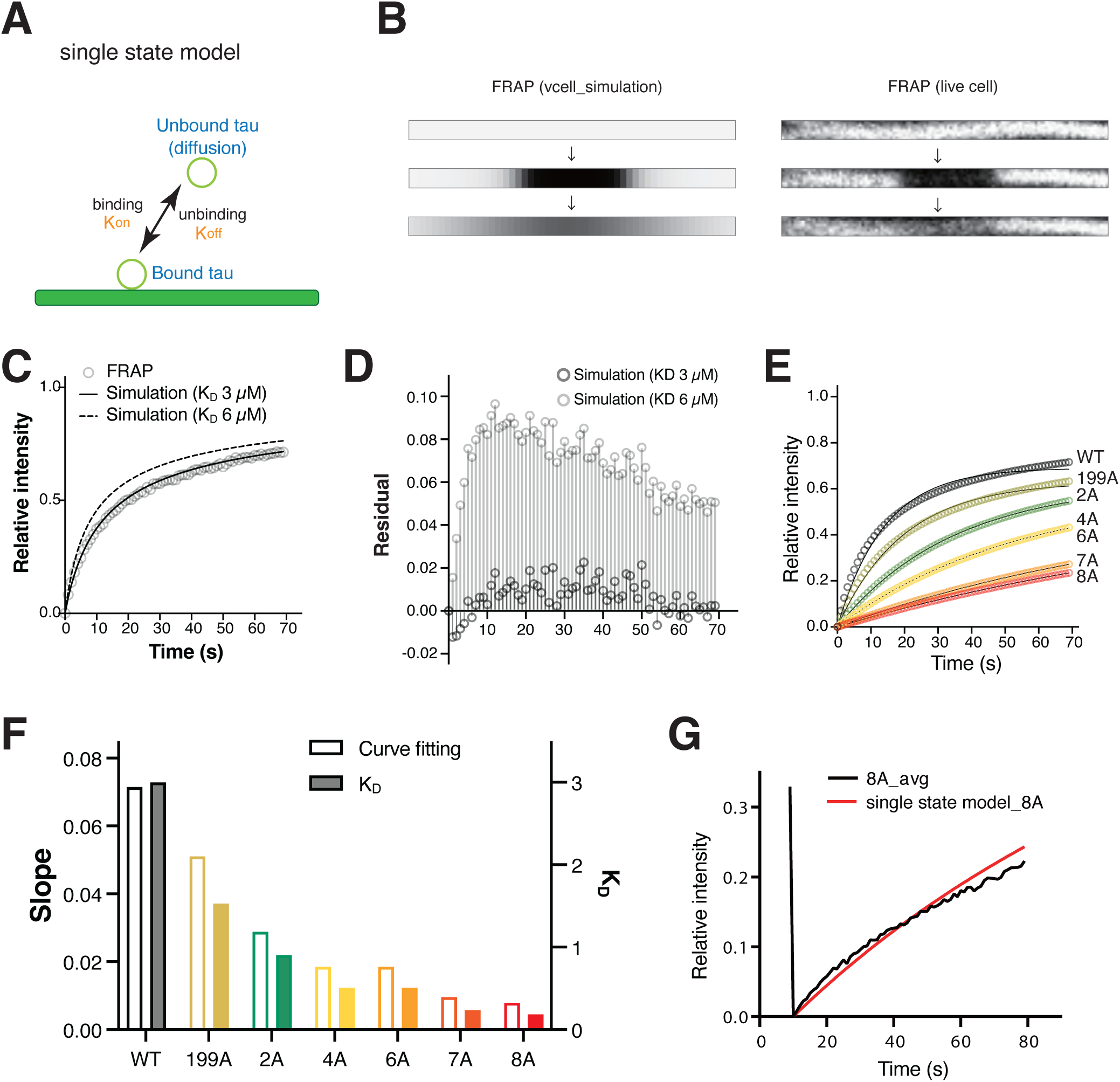
Simulation of FRAP experiments and estimation of binding kinetics of Tau to MTs. (A) Schematic of the simple "single-state model" used to describe Tau-MT interaction, with a single on-rate (Kon) and off-rate (Koff). (B) Comparison of simulated (left) and experimental (right) fluorescence images showing the bleached region before bleaching (top), immediately after bleaching (middle), and after recovery (bottom). (C) Fitting of the simulation curves to the experimental data for WT-Tau (gray line). Curves for K_D_= 3 µM (solid line) and 6 µM (dashed line) are shown. (D) Residual plot for the fit shown in (C) for K_D_= 3 µM (dark grey) and 6 µM (light grey). (E) Fitting of the simulation curves to the experimental data of the series of alanine mutants. (F) Quantitative analysis showing the FRAP recovery slope derived from the simulations (Curve fitting, white bars) and the dissociation constant (K_D_, colored bars) that provided the best fit for each mutant. The KD values progressively decrease as the number of alanine substitutions increases, indicating higher binding affinity. (G) Comparison of the experimental data for the 8A mutant (8A_avg, black line) and the best-fit curve from the single-state model (red line). A clear deviation is apparent, indicating the model is less successful in capturing the dynamics of the 8A mutant.

**Table 3.** Condition for the simulation (single state model)

| | $K_D$ |
| --- | --- |
| WT | 3 μM |
| 199A | 1.35 μM |
| 2A | 905 nM |
| 4A | 510 nM |
| 6A | 510 nM |
| 7A | 235 nM |
| 8A | 185 nM |

We then used the same model to analyze the alanine mutants and determined best fit values (Figure 4E). First, we reanalyzed the simulated curves with single exponential functions and found that they yielded similar slopes to those obtained with the experimental data and showed similar graded reduction with increasing number of alanine substitution (Figure 4F). K_D_ values which yielded the best fit to the data in our search were 1.35 µM (for 199A), 905 nM (for 2A), 510 nM (for 4A), 510 nM (for 6A), 235 nM (for 7A), and 185 nM (for 8A) (Figure 4F). This indicates that dephosphorylation of these residues in PRR2 gradually or additively enhances and stabilizes MT-binding of Tau, as suggested by the experimental data (see Figure 3).

However, it was less successful in capturing the dynamics of the mutants, where clear deviations between the simulated curve and the experimental data were apparent particularly for 8A (Figure 4G). To improve the model, particularly for the 8A mutant, we switched to a "two-state model" (Figure 5A). We have observed significant variability in FRAP data for WT-Tau and 8A-Tau, presumably due to several factors including variability in the phosphorylation states. Therefore, we implemented two Tau populations with distinct kinetic properties: a "fast-binding" (high K_D_) and a "slow-binding" (low K_D_) kinetics (Figure 5B; Table 4). To determine the higher- and lower-kinetic values, we plotted FRAP data from individual cell (Figure 5B) and simulated curves from the single state model with varying K_D_ values. From these plots, we made rough estimates of kinetic values for WT- and 8A-Tau for the two-state model. Then, the ratio of these two components providing best fit was searched using reciprocal and goodness of fit. This framework provided a visually superior fit for both WT-Tau and, most notably, for the 8A mutant (Figure 5C) with the kinetics values and the ratio of fast and slow populations shown in Table 4. We reckon two observations from this analysis: 1) both fast and slow kinetics became lower for 8A-Tau, and 2) the ratio of fast to slow components changed from 19:11 of WT to 1:3 of 8A. We have then analyzed the other mutants with this two-state model. Although this analysis is complicated in a way that it involves the two components and their ratios, the analyses clearly showed that both slow and fast K_D_ values were reduced with alanine substitution (Figure 5D; Table 4). Particularly, fast K_D_ values dropped dramatically even with a single 199A mutation (Figure 5D). Also, the proportion of the fast component showed clear trend of gradual decrease from WT to 8A.

**Figure 5.**
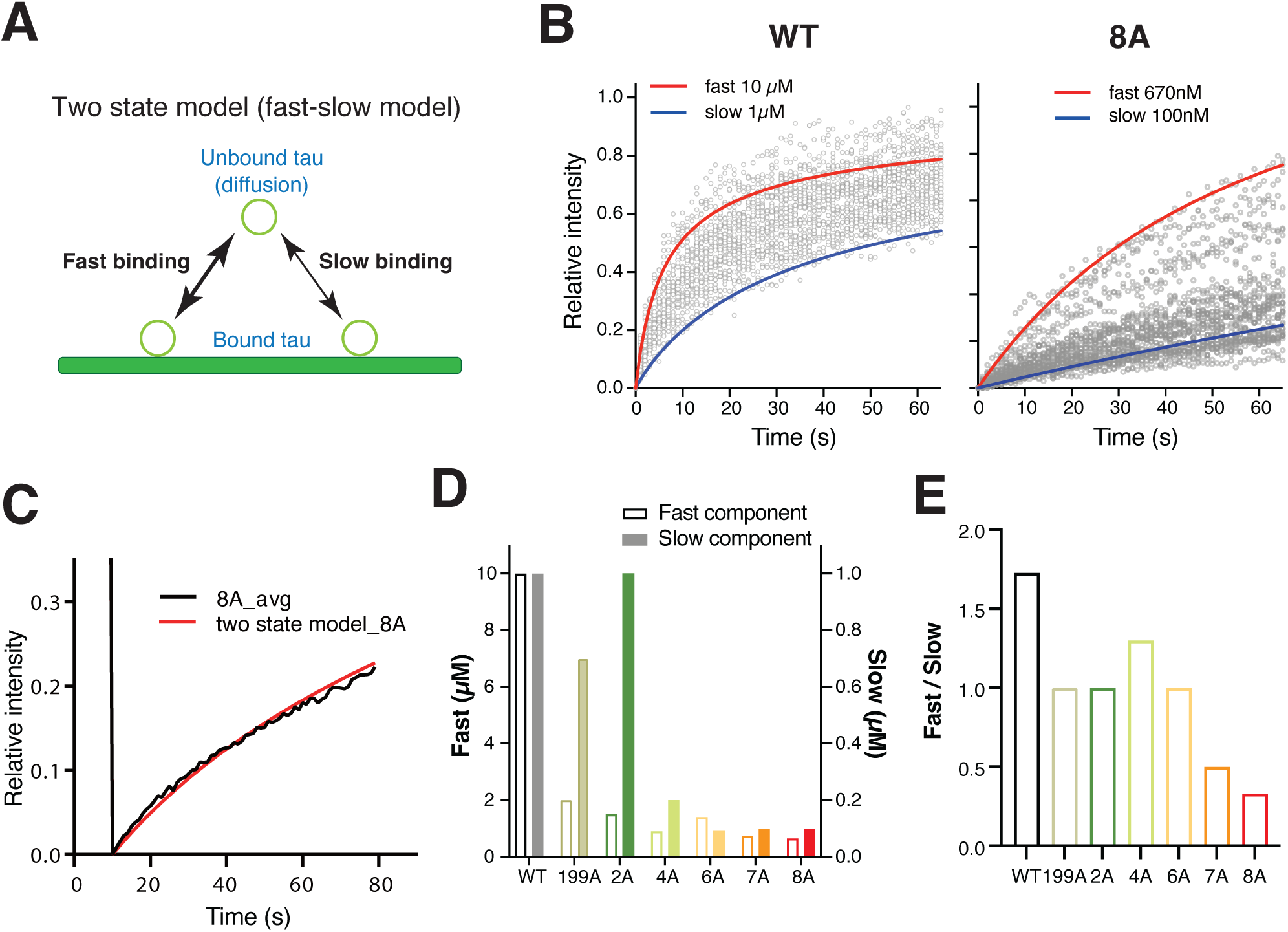
Refined simulation with the 2-state binding model. (A) Schematic of the "two-state model (fast-slow model)," which assumes two Tau populations with distinct kinetic properties. (B) FRAP data from individual cells (gray dots) for WT (left) and 8A (right). The overlaid curves approximate the upper ("fast") and lower ("slow") bounds of the data distribution, used to estimate the kinetic parameters for the two-state model. (C) Superior fit of the two-state model simulation (red line) to the experimental data for the 8A mutant (black line), showing a significant improvement over the single-state model (Fig. 4G). (D) Dissociation constants (K_D_) for the fast component (light bars) and slow component (dark bars) for each mutant, as determined by the model fitting. (E) The ratio of the fast to slow components for each mutant. A progressive shift towards the slow-binding population is observed as the number of alanine substitutions increases.

**Table 4.** Condition for the simulation (two state model/Fast-slow model)

|  | Fast | Slow | Fast/Slow ratio |
| --- | --- | --- | --- |
| WT | 10 μM | 1 μM | 19:11 |
| 199A | 3 μM | 700 nM | 2:1 |
| 2A | 1.5 μM | 1 μM | 1:1 |
| 4A | 900 nM | 200 nM | 17:13 |
| 6A | 1.4 μM | 90 nM | 1:1 |
| 7A | 750 nM | 100 nM | 1:2 |
| 8A | 670 nM | 100 nM | 1:3 |

Taken together, the computational analyses of the FRAP data supports the idea that the phosphorylation state of PRR2 gradually affect the binding kinetics of Tau to MTs in developing neurons.

## Discussion

In this study, we revealed a novel regulatory mechanism where the number of non-phosphorylated sites in Tau’s proline-rich region 2 (PRR2) acts as a ‘rheostat’ to fine-tune its binding to MTs using quantitative fluorescence imaging and computational approaches.

It is difficult to determine binding kinetics of MT-associated proteins in general and more so in living neurons. Here, we combined FRAP microscopy and computational modeling to gain reasonable estimates of binding kinetics of Tau-MT interaction. In particular, we reproduced FRAP experiments in silico with experimentally determined or literature-based constraints other than the binding kinetics (diffusion coefficients, and concentrations) and geometric settings similar to the experiments. Using these approaches, we found that WT-Tau exhibits an intracellular K_D_ of 3 µM in immature living neurons. This value is consistent with the wide range of K_D_ values (∼0.01 µM to ∼2 µM) reported in the literature, which vary significantly depending on experimental conditions and methods (Butner and Kirschner, 1991; Fauquant et al., 2011; Hervy and Bicout, 2019; Makrides et al., 2004). We could not determine K_on_ and K_off_ values since the binding reaction should be quite fast, and our numerical simulation is based on equilibrium. We were only able to examine that the lower bound of these kinetic parameters were 0.1 µM^-1^s^-1^ and 0.3 s^-1^, respectively. Surprisingly, the 8A dephospho-mimetic mutant exhibited a significantly lower K_D_ in the submicromolar range. These indicate that WT-Tau is highly phosphorylated in the PRR2 and probably in other regions in immature neurons used in the study. This is consistent with that Tau is generally highly phosphorylated in developing brain (Watanabe et al., 1993; Mandell and Banker, 1996).

Our findings, particularly the enhanced MT binding resulting from the dephospho-mimetic mutations in PRR2, are in good agreement with literatures (Kiris et al., 2011; Schwalbe et al., 2015). Furthermore, we extend this finding by revealing a previously uncharacterized aspect: by analyzing the series of alanine mutants, we show that the strength of this interaction is gradually modulated depending on the total number of phosphorylation sites. This suggests that Tau’s function may be precisely controlled, not by a single phosphorylation site, but by the overall level of phosphorylation across the entire PRR2 region. Another important finding is that the PRR2 domain regulates Tau’s MT binding independently of the MTBD. Our analysis of the 8AdelMTBD mutant shows that the phosphorylation state of PRR2 can alter Tau’s affinity for MTs even in the absence of the MTBD. This may be inconsistent with the CryoEM structure of Tau-MT complex, which did not exhibit obvious PRR-MT interaction (Brotzakis et al., 2021). However, we and others have shown that PRR2 does participate in MT-binding in vitro (Butner and Kirschner, 1991; Goode et al., 1997; Gustke et al., 1994). Furthermore, recent in vitro study showed that indeed PRR binds to MTs by itself and that its phosphorylation reduces the binding (Acosta et al., 2025). Therefore, the findings here and by others supports a more sophisticated model where the MTBD acts as a primary ‘anchor’, while the PRR2 domain functions as a tuner that dynamically modulates the strength of the interaction.

While our data indicate the role of PRR2 phosphorylation state on MT-binding, a complete understanding of Tau’s function requires placing this mechanism within the context of the entire protein. It is well-known that Tau is phosphorylated at numerous sites along its length. For instance, phosphorylation within the MTBD is known to directly detach Tau from MTs (Biernat et al., 1993; Schneider et al., 1999 Haj-Yahya et al., 2020), while phosphorylation in the N-terminal region has been suggested to alter Tau’s overall conformation. These effects might have been reflected in our FRAP data. The observed heterogeneity may not be merely random noise but may reflect the inherent biological variability in the post-translational modification state, particularly the phosphorylation state, of Tau. For WT-Tau, individual neurons likely maintain different equilibrium levels of endogenous phosphorylation in the PRR2 and other regions, which may result in the cell-to-cell variability of FRAP. For the 8A mutant, while the specific eight sites are "locked" in a dephospho-mimetic state, Therefore, the variability in 8A would reflect promiscuous phosphorylation states of other protein domains. Although we currently do not know the sites and extent of phosphorylation of endogenous and exogenous Tau in living neurons, with future development of single-cell phosphoproteome our study may serve as a foundation to study the roles of phosphorylation sites outside of PRR2. As the overall phosphorylation level of Tau is significantly altered under pathological conditions, understanding how these phosphorylation sites affect MT-binding and other aspects of Tau protein is of significance.

Several limitations to the present study should also be noted. Since we utilized alanine substitution as pseudo-dephosphorylation, we cannot rule out the structural or other unwanted effects of these mutations on MT-bindings. Since this type of mutation is widely used to study phosphorylation, and because we previously observed the opposite effect of pseudo-phosphorylation mutations (glutamate substitution) on MT-binding (Rodríguez-Martín et al., 2013), we are somewhat confident that the mutation mimics dephosphorylation, particularly together with the observed graded effects of alanine substitution. However, because the effect size of 8E-Tau was very small, we opted not to pursue individual or lesser number of E substitutions than 8E. As we have previously shown that FRAP of WT-Tau slightly slows down in more mature neurons (Iwata et al., 2019), using more mature neurons may provide a wider window of FRAP changes to probe pseudo-phosphorylation mutants in future considerations. Regarding this, we also rely on endogenous capability of phosphorylating unchanged residues to extract the effect of pseudo dephosphorylation on PRR2. In an in vitro binding assay, recombinant 8A-Tau would be indistinguishable from WT-Tau because they are not phosphorylated in E. Coli. Here, developing neurons can phosphorylate Tau at high levels (Watanabe et al., 1993), and thus we were able to detect the effect of the mutants. Another important limitation is the use of FRAP data for simulation. Because the temporal resolution is quite limited for our FRAP experiments, we were unable to determine how fast the binding reaction could be in the simulation. Also, although the model comparisons indicated that 8A results were better fitted by a model with at least two different pools, we cannot determine what these pools represent and whether two is sufficient. At least, this is consistent with the heterogeneity in individual FRAP curves observed presumably because of heterogeneous phosphorylation states of Tau in living neurons. Lastly, we did not implement fibrous nature of MTs in the model. This may be quite important, as it would increase local concentration of Tau binding sites on MTs and may facilitate re-binding of Tau. A model incorporating the fibrous nature MTs should provide more accurate picture of Tau binding in future studies.

## Acknowledgement

We thank Dr. Yoji Yonemura for expert advice on cell culture, cloning, and live-cell imaging techniques. We also thank Ms.Yuri Sakai for valuable technical assistance with the preparation of cultured neurons. This work was supported by the Sasakawa Scientific Research Grant from The Japan Science Society and JST SPRING (JPMJSP2129 to RN). Additional support was provided through the JSPS Core□to□Core Program A Advanced Research Networks (to HM) and JSPS KAKENHI Grant-in-Aid for Scientific Research(B) 23K24207 (to TM). The Virtual Cell is supported by NIH Grant R24 GM137787 from the National Institute for General Medical Sciences.

